# Evolving antibody evasion and receptor affinity of the Omicron BA.2.75 sublineage of SARS-CoV-2

**DOI:** 10.1101/2023.03.22.533805

**Authors:** Qian Wang, Zhiteng Li, Yicheng Guo, Ian A. Mellis, Sho Iketani, Michael Liu, Jian Yu, Riccardo Valdez, Adam S. Lauring, Zizhang Sheng, Aubree Gordon, Lihong Liu, David D. Ho

## Abstract

SARS-CoV-2 Omicron BA.2.75 has diversified into multiple subvariants with additional spike mutations, and several are expanding in prevalence, particularly CH.1.1 and BN.1. Here, we investigated the viral receptor affinities and neutralization evasion properties of major BA.2.75 subvariants actively circulating in different regions worldwide. We found two distinct evolutionary pathways and three newly identified mutations that shaped the virological features of these subvariants. One phenotypic group exhibited a discernible decrease in viral receptor affinities, but a noteworthy increase in resistance to antibody neutralization, as exemplified by CH.1.1, which is apparently as resistant as XBB.1.5. In contrast, a second group demonstrated a substantial increase in viral receptor affinity but only a moderate increase in antibody evasion, as exemplified by BN.1. We also observed that all prevalent SARS-CoV-2 variants in the circulation presently, except for BN.1, exhibit profound levels of antibody evasion, suggesting this is the dominant determinant of virus transmissibility today.

## Introduction

The Omicron variant of severe acute respiratory syndrome coronavirus 2 (SARS-CoV-2) continues to evolve, giving rise to several dominant subvariants worldwide. One particularly notable subvariant is designated as BA.2.75 (Wang et al., 2022d) **(Figure 1A)**. Since its detection in India in early May 2022, the Omicron BA.2.75 subvariant has rapidly spread to over 108 countries, competing with the predominant BQ and XBB subvariants, and its progenies are now responsible for over 8.54% of new SARS-CoV-2 cases worldwide **(Figure 1B)**. Instead of observing the emergence of a singular dominant form, the recently circulating BA.2.75-derived subvariants remain relatively genetically diverse and demonstrate different evolutionary pathways.

**Figure 1.**
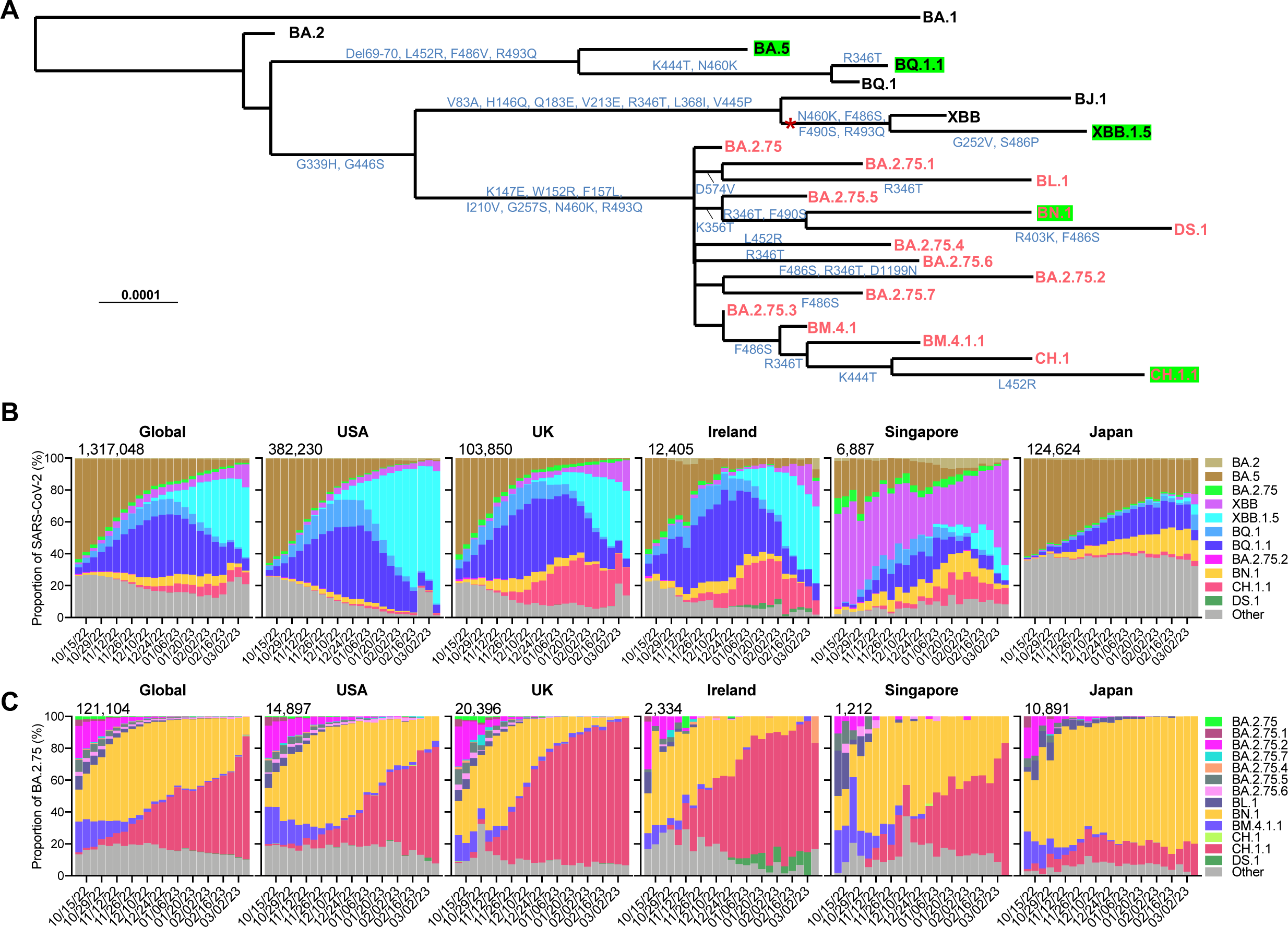
Spike alterations and prevalence of BA.2.75 subvariants. (**A**) Phylogenetic tree of selected BA.2.75 subvariants and current variants of concern (VOCs). The mutations on the branches showed the spike amino acid alterations of each variant. The recombination event for XBB from BJ.1 and BA.2.75 is denoted by the asterisk. The BA.2.75 subvariants are highlighted in red, and green boxes indicate current globally dominant variants with a frequency over 2%. Proportions of SARS-CoV-2 VOCs (**B**) and frequencies of BA.2.75 subvariants among BA.2.75 (**C**) in GISAID from October 2022 to March 2023. The cumulative number of sequences in the denoted time period is displayed at the upper right corner of each graph. See also **Figure S1**.

The current most frequently observed subvariants of BA.2.75 are CH.1.1, which has additional R346T, K444T, L452R, and F486S mutations, and BN.1, which has additional R346T, K356T, and F490S mutations, in their respective spikes **(Figure 1A)**. Globally, CH.1.1 and BN.1 account for 77.2% and 11.1% of recent infections by BA.2.75 progenies, respectively. Interestingly, CH.1.1 appears to be more dominant among BA.2.75 subvariants in European and American countries, including the USA, the UK, and Ireland, while BN.1 is more dominant among BA.2.75 infections in Asian countries, such as Japan **(Figure 1C)**. Another BA.2.75 subvariant drawing attention is DS.1, which has additional R403K and F486S mutations beyond BN.1 and was rapidly rising in Ireland weeks ago, where XBB.1.5, BQ.1.1, and CH.1.1 co-circulate. Other subvariants derived from BA.2.75 are also noteworthy, as they also carry spike mutations, including R346T, K444T/M, L452R, F486S, or F490S **(Figure 1A)**, which have been reported to impair monoclonal antibody (mAb) or polyclonal serum neutralization (Cao et al., 2023; Starr et al., 2022; Wang et al., 2022c; Wang et al., 2022e).

The spike proteins of the Omicron BA.2.75 sublineage feature multiple convergent mutations that were previously observed in other Omicron variants, such as R346T, K444T, L452R, F486S, and F490S (Wang et al., 2022c; Wang et al., 2023; Wang et al., 2022e), as well as three newly identified mutations: R403K, K356T, and D574V (**Figure S1**). The expansion of these spike mutations observed in the new BA.2.75 subvariants therefore raise concerns about their impact on the effectiveness of current vaccines and antibody therapies. Here, our study addresses this concern and provides additional insight into SARS-CoV-2 evolutionary trajectory.

## Results

### Divergent receptor-binding affinities

Viral entry into the cell begins with binding to a receptor. Therefore, transmission advantages of BA.2.75 subvariants may be associated, in part, with their binding affinity to the relevant viral receptor. Here, we measured the binding affinity between human angiotensin converting enzyme 2 (hACE2) and each spike of several major BA.2.75 subvariants, as well as other important Omicron subvariants and viruses with select single mutations, by surface plasmon resonance (SPR) **(Figure 2)**. Overall, BA.4/5, BF.7, BQ.1, BQ.1.1 and XBB.1.5 exhibited higher binding affinity to the receptor compared to the D614G strain, as we previously observed (Wang et al., 2023; Wang et al., 2022e). Among BA.2.75 subvariants, several displayed lower binding affinities compared to parental BA.2.75 following the acquisition of more spike mutations, including CH.1, CH.1.1 and DS.1 **(Figure 2A)**. Interestingly, when BA.2.75 carries an additional mutation R346T (BA.2.75.6), K356T (BA.2.75.5), L452R (BA.2.75.4), D574V (BA.2.75.1), or R403K, the hACE2-binding affinity was enhanced, even though these mutations are not in direct contact with the binding interface (**Figures 2B and 2C**). In contrast, other single mutations (K444M, K444T, F490S, and D1199N) did not dramatically alter binding affinity. Similar to F486V found in BA.4/5, F486S was also shown to greatly reduce binding affinity (Wang et al., 2022c). Remarkably, BN.1, likely through the combination of R346T and K356T, had the highest binding affinity among BA.2.75 subvariants tested (4.0-fold that of BA.2.75). In short, the receptor binding affinity for BA.2.75 subvariants has evolved in two distinct directions: one exhibiting increased affinity compared with BA.2.75, as observed in BN.1, while another demonstrating slightly decreased affinity, as observed for CH.1.1 and DS.1 **(Figure 2C)**.

**Figure 2.**
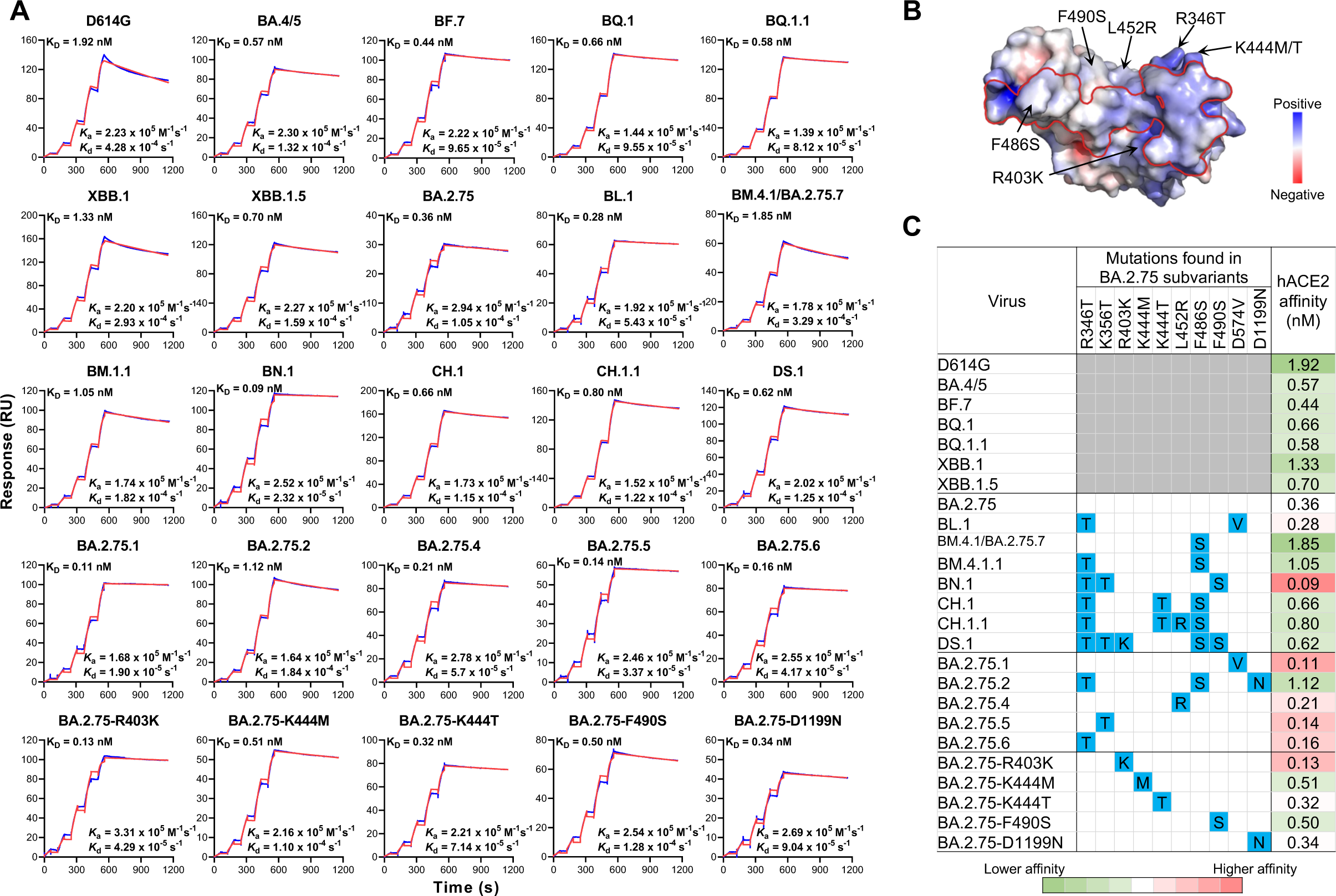
Binding affinities of SARS-CoV-2 spike proteins with human angiotensin converting enzyme 2 (hACE2) as measured by SPR. **(A)** Characterization of the binding between spike proteins with hACE2 tested by SPR. The raw and fitted curves are represented by blue and red lines, respectively. **(B)** The electrostatic surface potential of the RBD in top view, with red and blue corresponding to negative and positive charges, respectively. The red line on the RBD surface indicates the footprint of ACE2. Black arrows indicate the surrounding mutations found in BA.2.75 subvariants. **(C)** The summarized profile of viral receptor affinities of spike proteins. Spike mutations found in each of the indicated subvariants in addition to BA.2.75 are highlighted in blue. Enhanced ACE2 affinities compared with that of BA.2.75 are highlighted in red, while reduced affinities are in green. The results shown are representative of those obtained in two independent experiments.

### Evasion of neutralization by monoclonal antibodies

To investigate the antibody evasion properties of BA.2.75 sublineage, we generated vesicular stomatitis virus (VSV)-pseudotyped viruses of each subvariant and the R403K, K444M, K444T, F490S, and D1199N point mutants in the background of BA.2.75 (denoted BA.2.75-R403K, BA.2.75-K444M, BA.2.75-K444T, BA.2.75-F490S, and BA.2.75-D1199N, respectively) **(Figure 3A)**. We then assessed their neutralization profile to a panel of 30 mAbs that had retained good potency against both D614G and BA.2.75 parental virus by targeting multiple epitopes on the viral spike. Among these mAbs, 27 were directed to the four epitope classes in the receptor binding domain (RBD), including Brii-196 (amubarvimab) (Ju et al., 2020), Omi-3 (Nutalai et al., 2022), Omi-18 (Nutalai et al., 2022), BD-515 (Cao et al., 2021), COVOX-222 (Dejnirattisai et al., 2021), XGv051 (Wang et al., 2022b), XGv347 (Wang et al., 2022a), ZCB11 (Zhou et al., 2022), S2E12 (Tortorici et al., 2020), COV2-2196 (tixagevimab) (Zost et al., 2020), LY-CoV1404 (bebtelovimab) (Westendorf et al., 2022), 2-7 (Liu et al., 2020), XGv289 (Wang et al., 2022a), XGv264 (Wang et al., 2022b), S309 (sotrovimab) (Pinto et al., 2020), P2G3 (Fenwick et al., 2022), SP1-77 (Luo et al., 2022), BD55-5840 (Cao et al., 2022), BD55-3152 (Cao et al., 2022), XGv282 (Wang et al., 2022a), BD-804 (Du et al., 2021), A19-46.1 (Wang et al., 2021), 35B5 (Wang et al., 2022g), JMB2002 (Yin et al., 2022), Brii-198 (romlusevimab) (Ju et al., 2020), COV2-2130 (cilgavimab) (Zost et al., 2020), and 10-40 (Liu et al., 2022). The other three mAbs, C1520 (Wang et al., 2022h), C1717 (Wang et al., 2022h), and S3H3 (Hong et al., 2022) target the N-terminal domain (NTD), NTD-SD2 (subdomain 2), and SD1 (subdomain 1), respectively. In **Figure 3B**, the footprints of those mAbs with structural information available were drawn on the spike or the RBD with the key mutations found in the BA.2.75 subvariants highlighted.

**Figure 3.**
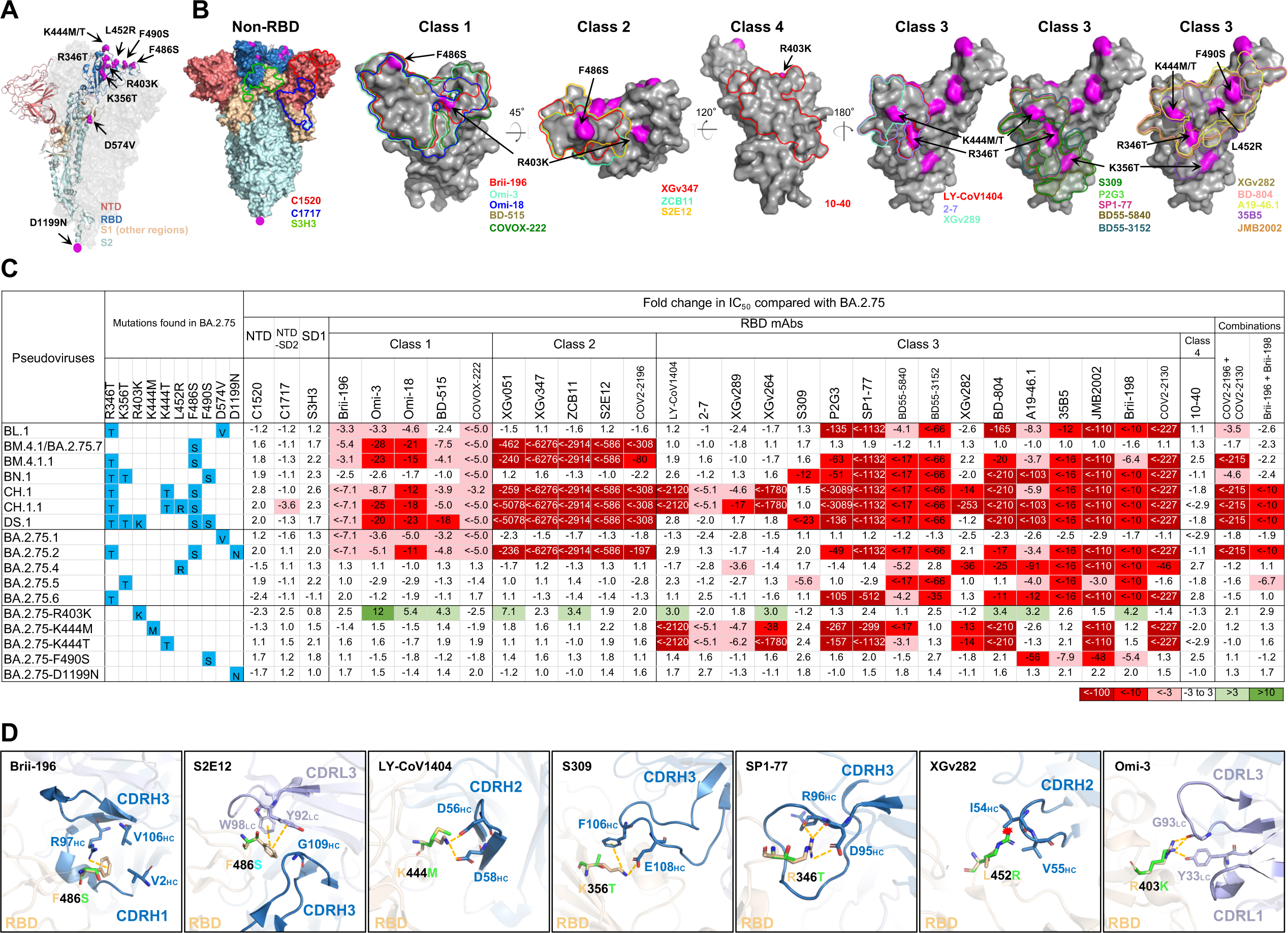
Resistance of pseudotyped BA.2.75 subvariants to neutralization by monoclonal antibodies (mAbs). **(A)** Key spike mutations found in BA.2.75 subvariants. Mutations are highlighted in magenta. **(B)** Footprints of NTD-, NTD-SD2- and SD1-directed neutralizing mAbs on spike, and RBD class 1 to class 4-directed neutralizing mAbs on RBD. Mutations found in BA.2.75 subvariants are highlighted in magenta. **(C)** Fold change in IC_50_ values of BA.2.75 subvariants relative to BA.2.75, with resistance to neutralization highlighted in red and sensitization in green. Spike mutations found in each of the indicated subvariants in addition to BA.2.75 are highlighted in blue. **(D)** Structural modeling of the impact on mAbs for the F486S, K444M, K346T, K356T, L452R, and R403K mutations. Clash is shown as the red asterisk; the interactions are shown as yellow dashed lines. See also **Table S1**.

The raw IC_50_ (the 50% inhibitory concentration) values for each mAb against each pseudovirus are summarized in **Table S1**, and the fold changes in IC_50_ values compared to that of BA.2.75 are presented in **Figure 3C**. Overall, the non-RBD mAbs and class 4 RBD mAb (C1520, C1717, S3H3, and 10-40) generally did not have impaired neutralization activity against the BA.2.75 subvariants, with the exception of C1717, which had a 3.6-fold drop in neutralizing CH.1.1. Class 1, 2, and 3 RBD mAbs exhibited diverse neutralization profiles. BM.4.1 and BA.2.75.7 (denoted as BM.4.1/BA.2.75.7 as they share an identical spike) partially or completely escaped neutralization by both class 1 and class 2 RBD mAbs because of the F486S mutation. BA.2.75.1 resisted RBD class 1 mAbs, due to the D574V mutation. BA.2.75.4, BA.2.75.5, and BA.2.75.6 demonstrated substantial resistance to some of the RBD class 3 mAbs after acquiring L452R, K356T, and R346T, respectively. The single mutation D1199N found in BA.2.75.2 did not alter the neutralization profile of BA.2.75, while the mutations K444M/T and F490S both impaired the neutralizing activity of some class 3 RBD mAbs.

BN.1 also exhibited relative resistance to some class 3 RBD mAbs, due to R346T, K356T, and F490S. The most strikingly resistant of the subvariants were BM.4.1.1, CH.1, CH.1.1, DS.1, and BA.2.75.2, which showed neutralization resistance to class 1 and 2 RBD mAbs and most class 3 RBD mAbs. CH.1 and CH.1.1 were most evasive to neutralization by this panel of mAbs, as only 2 of 27 RBD-directed mAbs retained unchanged potency against these subvariants, followed by DS.1, which impaired neutralization by 21 of 27 RBD antibodies. Surprisingly, we observed that a mutation unique to DS.1, R403K, in fact slightly sensitized BA.2.75 to neutralization by 10 of 26 class 1, 2, and 3 RBD mAbs tested **(Figure 3C)**.

We also studied several clinically authorized antibodies and cocktails against BA.2.75 subvariants, including COV2-2130 and COV2-2196 (also known as Evusheld), the combination of Brii-196 (amubarvimab) and Brii-198 (romlusevimab), and LY-CoV1404 (bebtelovimab). Evusheld was rendered inactive or greatly impaired against BA.2.75.2, BM.4.1.1, CH.1, CH.1.1, and DS.1 which all have the R346T mutation paired with F486S **(Figure 3C)**. Brii-196 + Brii-198, which was already greatly impaired against BA.2.75, further lost activity against BA.2.75.2, BA.2.75.5, CH.1, CH.1.1, and DS.1. Bebtelovimab was knocked out by the BA.2.75 single mutants carrying the K444M/T mutations, as well as by CH.1 and CH.1.1.

### Structural modeling of mAb-binding impairment in BA.2.75 subvariants

We conducted structural modeling to further investigate how mutations in the circulating BA.2.75 subvariants confer resistance or sensitization to mAbs against different epitopes **(Figure 3D)**. One of these mutations, F486S, disrupted a common cation-π interaction with R97 of Brii-196 from VH3-53 gene class (Yuan et al., 2020), as well as the interactions with Y92 and W98 for S2E12. The other RBD mutations, R346T, K356T, K444M/T, L452R, and F490S, are located on the outer surface of the RBD and within the epitope cluster of class 3 mAbs, which likely explains their loss of neutralizing activity. Specifically, the K444M and K444T mutations abolished two salt bridges interacting with D56 and D58 in CDRH2 of LY-CoV1404 (Wang et al., 2023), and the K356T mutation weakened S309 by breaking the salt bridge and cation-π interaction. Additionally, the K356T mutation may introduce an N-glycan at N354 in RBD, which would reduce the accessibility of the surrounding residues, thereby conferring a degree of resistance to the RBD class 3 mAbs. R346T, L452R, and F490S mutations have previously been observed in BA.4.6 (Jian et al., 2022; Wang et al., 2022e), BA.4/5 (Cao et al., 2022; Wang et al., 2022c), and Lambda (Kimura et al., 2022; Wang et al., 2022f), respectively. The R346T mutation removed the salt bridge and several hydrogen bonds with D95 and R96 in SP1-77, the L452R mutation created steric hindrance with I54 in XGv282, and F490S disturbed the cation-π interaction with R74 in XGv282 (Wang et al., 2023) **(Figure 3D)**. R403K mutation, a novel substitution sensitizing BA.2.75 to some RBD-directed mAbs, could retain the interaction with G93, and form an extra hydrogen bond with Y33 in Omi-3 **(Figure 3D)**.

### Enhanced evasion of serum neutralization

Given the increased evasion of BA.2.75 subvariants to mAb neutralization and the structural changes within multiple key epitopes, we next asked whether these subvariants were also capable of evading neutralization by sera from humans with prior immunity to SARS-CoV-2. We measured the neutralization resistance profiles of the BA.2.75 subvariants to sera from four different clinical cohorts: individuals who had received three doses of the wildtype mRNA vaccines (“3 shots WT”), three doses of the wildtype mRNA vaccines followed by one shot of the bivalent mRNA vaccines (“3 shots WT + bivalent”), and patients who had a BA.2 or BA.4/5 breakthrough infection after vaccination (“BA.2” and “BA.4/5 breakthrough”, respectively; **Table S2**). Their neutralization ID_50_ titers (50% inhibitory dilution) against D614G, BA.4/5, BA.2.75, and BA.2.75 subvariants are presented in **Figure 4A**. Consistent with our previous findings, BA.2.75 was 7.5-fold more resistant to the “3 shots WT” sera neutralization, while BA.4/5 was 1.8-fold more resistant than BA.2.75 (Wang et al., 2022d). The neutralization ID_50_ titers of this cohort were significantly lower against the new BA.2.75 subvariants than against BA.2.75, with the exception of BA.2.75.5. CH.1.1 showed the biggest drop in susceptibility to neutralization, of 11.9-fold, in the “3 shots WT” cohort. In addition, BL.1, BM.4.1/BA.2.75.7, BM.4.1.1, BN.1, CH.1, CH.1.1, and DS.1 significantly impaired the neutralization potency of the boosted sera by 2.1- to 11.0-fold, among which, BM.4.1.1, CH.1, CH.1.1, and DS.1 were more neutralization evasive than BA.2.75.2. The D1199N single mutation, as well as the K444M, K444T, and F490S mutations, did not strongly alter the neutralization resistance of BA.2.75 to the “3 shots WT” sera (0.9- to 1.4-fold changes in ID_50_ titers). Strikingly, concordant with increased mAb neutralization, R403K sensitized BA.2.75 to serum neutralization by 1.7-fold.

**Figure 4.**
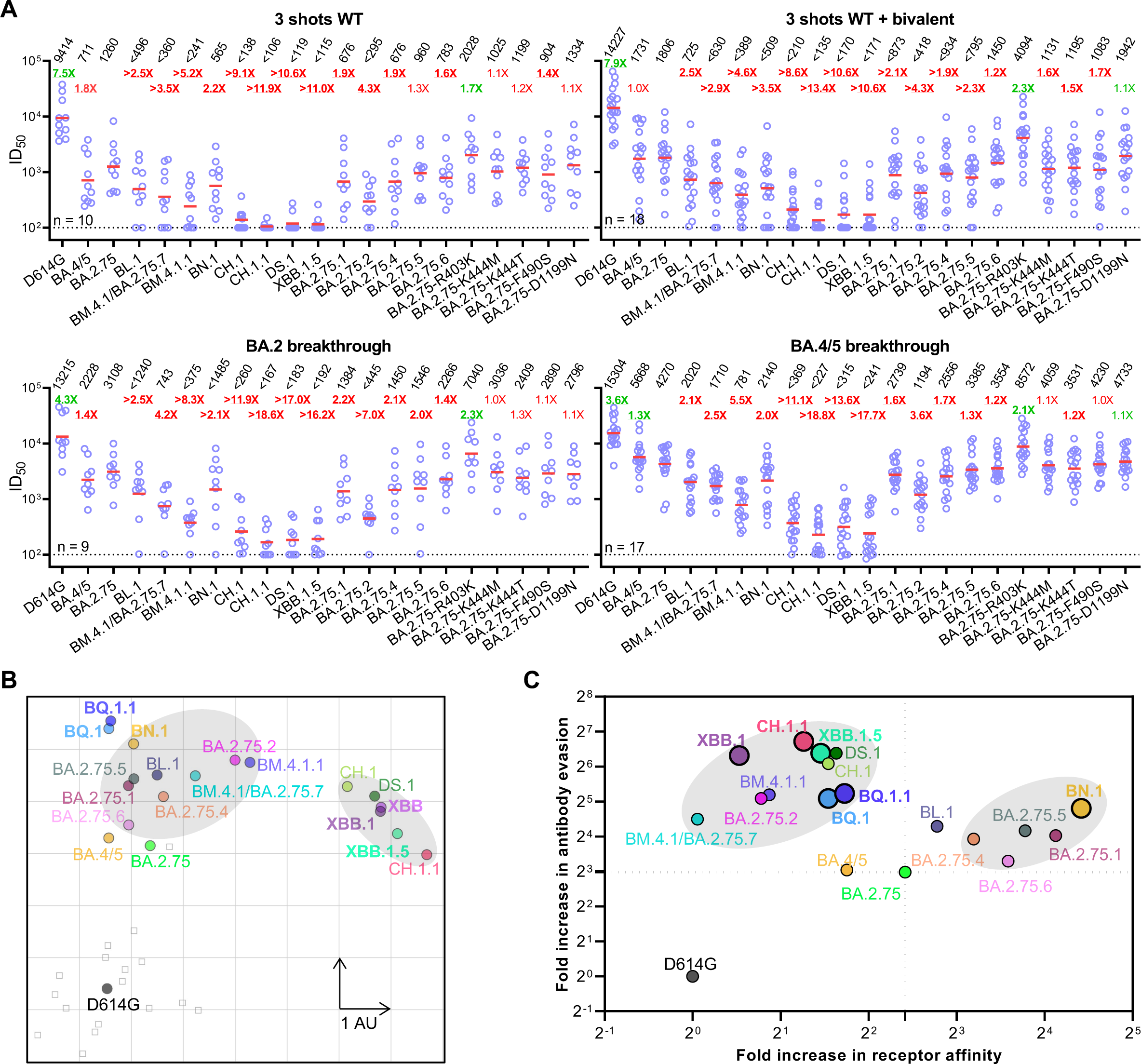
Neutralization of pseudotyped BA.2.75 subvariants by polyclonal sera from four clinical cohorts. **(A)** Neutralization of pseudotyped D614G and Omicron subvariants by sera from four different clinical cohorts. “3 shots WT” refers to individuals who received three doses of a COVID-19 WT mRNA vaccine, “3 shots WT + bivalent” refers to individuals vaccinated with three doses of the wildtype mRNA vaccine and subsequently one dose of a WA1/BA.5 bivalent mRNA vaccine, and breakthrough refers to individuals who received COVID-19 vaccines and were infected. The results are representative of those obtained in two independent experiments and shown as dots with geometric mean (red line). Values above the dots denote the raw geometric mean ID_50_ values and the sample size (n) for each group is shown on the lower left. The limit of detection is 100 (dotted line). Comparisons were made against BA.2.75 and the fold changes in ID_50_ values are shown, with resistance to neutralization highlighted in red and sensitization in green. Statistically significant fold changes (*p* <0.05, determined by using two-tailed Wilcoxon matched-pairs signed-rank tests) are highlighted in bold. **(B)** Antigenic map based on the neutralization data of “3 shots WT + bivalent” vaccinee sera. SARS-CoV-2 variants are shown as colored circles and sera are shown as gray squares. The x and y axes represent antigenic units (AU) with one unit corresponding to a two-fold serum dilution of the neutralization titer. **(C)** Changes to receptor-binding affinity and antibody evasion of Omicron BA.2.75 subvariants. The x-axis illustrates the fold change in ACE2-binding affinity of the Omicron subvariants relative to the D614G strain. The y-axis represents the relative immune evasion capability of Omicron subvariants in comparison to the D614G strain (fold change in geometric mean ID_50_ over “3 shots WT + bivalent” cohort). Black dashed lines correspond to equivalence to BA.2.75. See also **Table S2**.

A similar trend was also observed for the 3 shots WT + bivalent cohort and the BA.2 and BA.4/5 breakthrough cohorts. BL.1, BM.4.1/BA.2.75.7, BM.4.1.1, BN.1, and BA.2.75.2 impaired the neutralization potency of sera moderately more than BA.2.75, while CH.1, CH.1.1, and DS.1 exhibited substantially stronger antibody evasion to serum neutralization, similar to the current predominant Omicron subvariant XBB.1.5.

To visualize the antigenic relationship of the BA.2.75 subvariants, we used antigenic cartography (Smith et al., 2004) of the bivalent vaccine boosted serum neutralization results to construct a graphical map to display the antigenic distances among D614G, BA.4/5, XBB, XBB.1, XBB.1.5, BQ.1, BQ.1.1, and BA.2.75 subvariants (**Figures S2 and 4B**). In this rendering, each antigenic unit (AU) of distance in any direction corresponds to a two-fold change in ID_50_ titer. BA.2.75 and BA.4/5 displayed a similar antigenic distance to the bivalent vaccine boosted sera. The point mutations, R346T in BA.2.75.6, K356T in BA.2.75.5, L452R in BA.2.75.4, F486S in BM.4.1/BA.2.75.7, and D574V in BA.2.75.1, each increased the antigenic distance from the boosted sera compared to parental BA.2.75 by about 0.3, 1.14, 0,92, 1.47, and 1.02 AU, respectively, suggesting their importance in mediating resistance to polyclonal antibody neutralization. The combination of R346T, F486S, and D1199N (BA.2.75.2) was an average of 5.07 AU from bivalent sera, which is 1.96 AU further away than BA.4/5, suggesting an advantage of BA.2.75.2 over BA.4/5 in evading serum antibodies. BN.1 and BL.1 were 4.8 and 4.29 AU from boosted sera, respectively, which is at least 0.44 AU closer than BQ.1 and BQ.1.1, while BM.4.1.1 exhibited a similar distance to that of BQ.1 and BQ.1.1. Interestingly, CH.1, CH.1.1 and DS.1 were 6.09, 6.75 and 6.40 AU from bivalent vaccine boosted sera, which is similar to the 6.39 AU for XBB.1.5, illustrating the competitive resistance advantage of CH.1.1, DS.1 and XBB.1.5. Finally, we note that these BA.2.75 subvariants have diverged into two separate antigenic clusters, one set (e.g., BN.1 and BA.2.75.1) grouping with BQ subvariants and another (e.g., CH.1.1 and DS.1) grouping with XBB subvariants **(Figure 4B)**. These two clusters are antigenically quite distinct, highlighting the significant antigenic differences between these viruses.

### Evolution of BA.2.75 subvariant phenotypes

Lastly, we sought to examine whether there were common trends in receptor-binding and antibody evasion phenotypes across the BA.2.75 sublineage, by plotting the fold increases in hACE2-binding affinity versus the fold increases in antibody evasion to sera from the bivalent boosted cohort **(Figure 4C)**. Two general phenotypic combinations were evident: one group including CH.1.1 and DS.1 with substantially higher neutralization resistance but decreased ACE2 affinity, and another group including BN.1 with substantially higher ACE2 affinity but only moderately increased neutralization resistance. Placed in the context of other Omicron subvariants (e.g., XBB.1.5 and BQ.1.1), it is quite apparent that viral strains that are currently dominant in the circulation are primarily those with highest antibody evasion properties, again indicating this particular phenotype as the principal determinant of transmissibility in the population today **(Figure 4C)**.

## Discussion

To gain a deeper understanding of the evolution of the emerging Omicron BA.2.75 sublineage, we systematically evaluated antigenic and viral receptor-binding properties of major subvariants within this sublineage. Our experimental and *in silico* analyses revealed several critical mutations, including three not previously described (K356T, R403K, and D574V) (**Figure S1**), which conferred different degrees of antibody resistance and receptor-binding affinity (**Figures 2, 3, and 4**). Five RBD mutations, R346T, K356T, K444M/T, and F490S impaired the neutralization activities of some class 3 RBD mAbs, F486S led to a large loss of neutralizing activities of classes 1 and 2 RBD mAbs, and D574V conferred resistance to all of the class 1 RBD mAbs tested. Notably, the BA.2.75 subvariants with K444M/T and R346T paired with F486S evaded authorized antibodies bebtelovimab and Evusheld, respectively, which poses a new threat to individuals who need them therapeutically or prophylactically. Interestingly, we made the novel observation that R403K sensitizes BA.2.75 to neutralization by some class 1, 2, and 3 RBD mAbs **(Figure 3)**, while it substantially increases the receptor affinity. This affinity increase could be a compensatory mechanism to regain the fitness loss in receptor binding caused by mutations at F486 in the DS.1 subvariant, mechanistically similar to the R493Q mutation observed in BA.4/5 (Wang et al., 2022c).

In addition to R403K, our study shows that R346T, K356T, and D574V not only contributed to neutralization profile changes but also enhanced receptor-binding affinity, which sheds light on the continued co-evolution of immune evasion and factors affecting transmissibility of the virus (**Figures 2, 3, and 4**). This higher receptor binding affinity could potentially compensate for lower antibody evasion properties and allow for expansion, as exemplified by BN.1.

Overall, our investigations have shown that the evolutionary trajectory of BA.2.75 subvariants has diverged in two different directions: substantially higher neutralization resistance but slightly reduced ACE2 affinity, as seen in CH.1.1 and DS.1, and substantially higher ACE2-affinity but only moderately increased neutralization resistance, as seen in BN.1 **(Figure 4C)**. Globally, and particularly in the US, UK, Ireland and Singapore, BN.1 is being outcompeted by CH.1.1 **(Fig. 1C)**, although it remains unclear why the same has yet to occur in some Asian countries such as Japan.

BA.2.75 subvariants CH.1, CH.1.1, and DS.1 now rival XBB.1.5 in their resistance to monoclonal antibodies **(Figure 3C)** and polyclonal sera **(Figure 4A)**. XBB.1.5 is slightly more antibody evasive than BQ.1 and BQ.1.1 (Wang et al., 2023). It appears that highest level of resistance to antibody neutralization has been achieved by three Omicron sublineages (XBB.1/XBB.1.5, BQ.1/BQ.1.1, and CH.1.1), each utilizing a distinct mutational pathway to converge on a common phenotype **(Figure 1A)**. Seemingly, the most dominant SARS-CoV-2 strains presently are also most evasive to antibody neutralization **(Figure 4C)**. This correlation suggests that the ability to escape from antibody pressure is perhaps the dominant determinant of transmissibility in the population today.

## STAR METHODS

### Key resource table Resource availability

- Lead contact
- Materials availability
- Data and code availability

### Experimental model and subjects

- Sample collection
- Cell lines

### Method details

- Plasmids
- Protein expression and purification
- Surface plasmon resonance (SPR)
- Pseudovirus production
- Pseudovirus neutralization assay
- Phylogenetic analysis
- Antibody footprint and mutagenesis analysis
- Antigenic cartography

**QUANTIFICATION AND STATISTICAL ANALYSIS**

## ACKNOWLEDGEMENTS

This study was supported financially by the SARS-CoV-2 Assessment of Viral Evolution Program, NIAID, NIH (Subcontract No. 0258-A709-4609 under Federal Contract No. 75N93021C00014) and the Gates Foundation (Project No. INV019355) awarded to D.D.H., and through the National Institutes of Health Collaborative Influenza Vaccine Innovation Center (75N93019C00051) awarded to A.G.. We acknowledge Michael T. Yin and Magdalena E. Sobieszczyk for providing serum samples. We thank Carmen Gherasim, David Manthei, Emily Stoneman, Victoria Blanc, Pamela Bennett-Baker, Savanna Sneeringer, Lauren Warsinske, Theresa Kowalski-Dobson, Alyssa Meyers, Zijin Chu, Hailey Kuiken, Lonnie Barnes, Ashley Eckard, Kathleen Lindsey, Dawson Davis, Aaron Rico, Casey Juntila, Daniel Raymond, Mayurika Patel, Nivea Vydiswaran, Julie Gilbert, William J. Fitzsimmons from the IASO study team for providing serum samples. We thank all who contributed their data to GISAID.

## AUTHOR CONTRIBUTIONS

L.L. and D.D.H. conceived this project. Q.W. and L.L. constructed the spike expression plasmids. Q.W., I.A.M., and L.L. conducted pseudovirus neutralization assays. Q.W., Z.L. and L.L. purified SARS-CoV-2 spike proteins, performed SPR assay and conducted structural analyses. Y.G. and Z.S. conducted bioinformatic analyses and generated antigenic map. Q.W. managed the project. J.Y. and M.L. expressed and purified antibodies. R.V., A.S.L., and A.G. provided clinical samples. L.L. and D.D.H. directed and supervised the project. Q.W., Z.L., Y.G., I.A.M., S.I., L.L., and D.D.H. analyzed the results and wrote the manuscript.

## STAR METHODS

### Key resource table

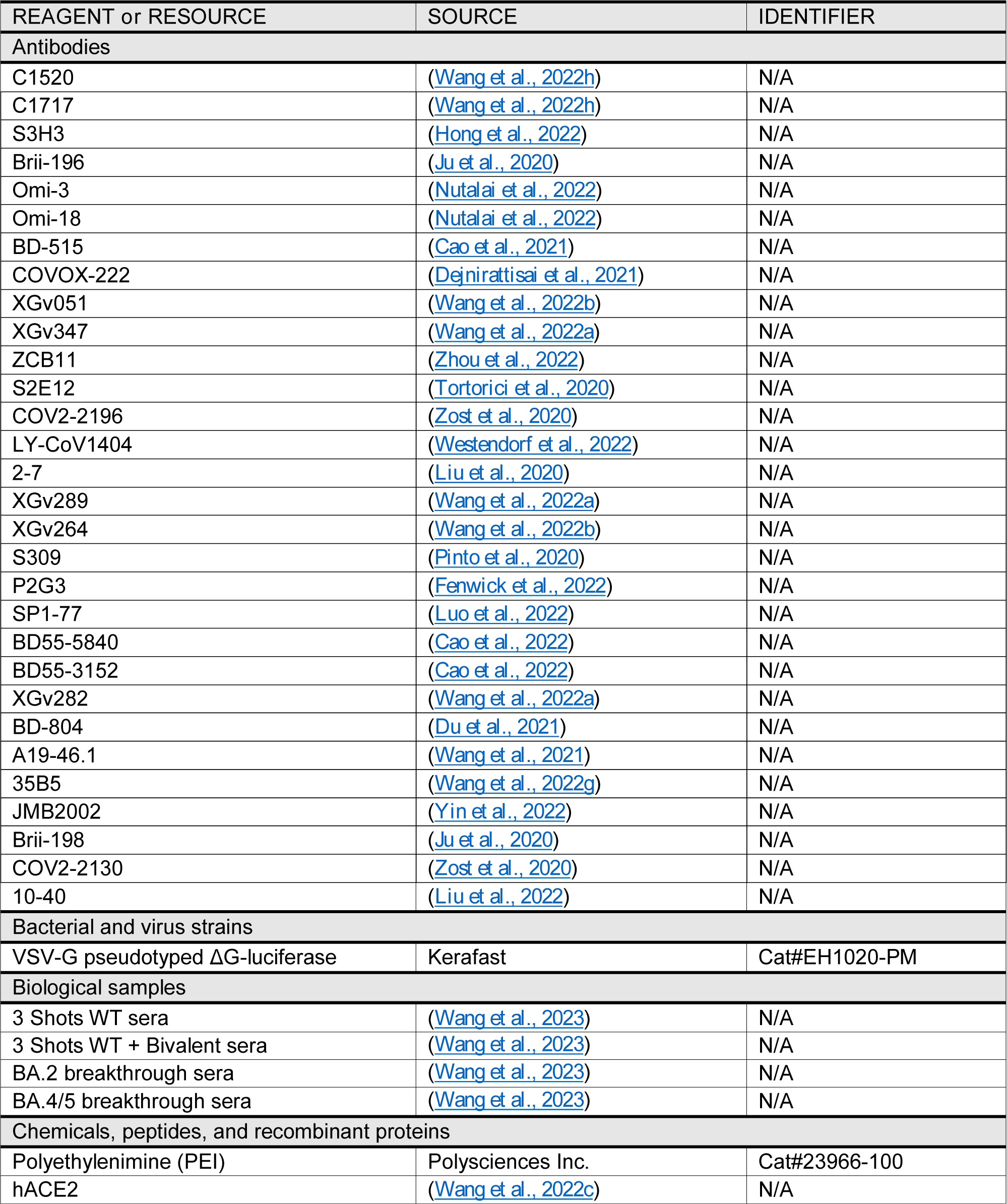
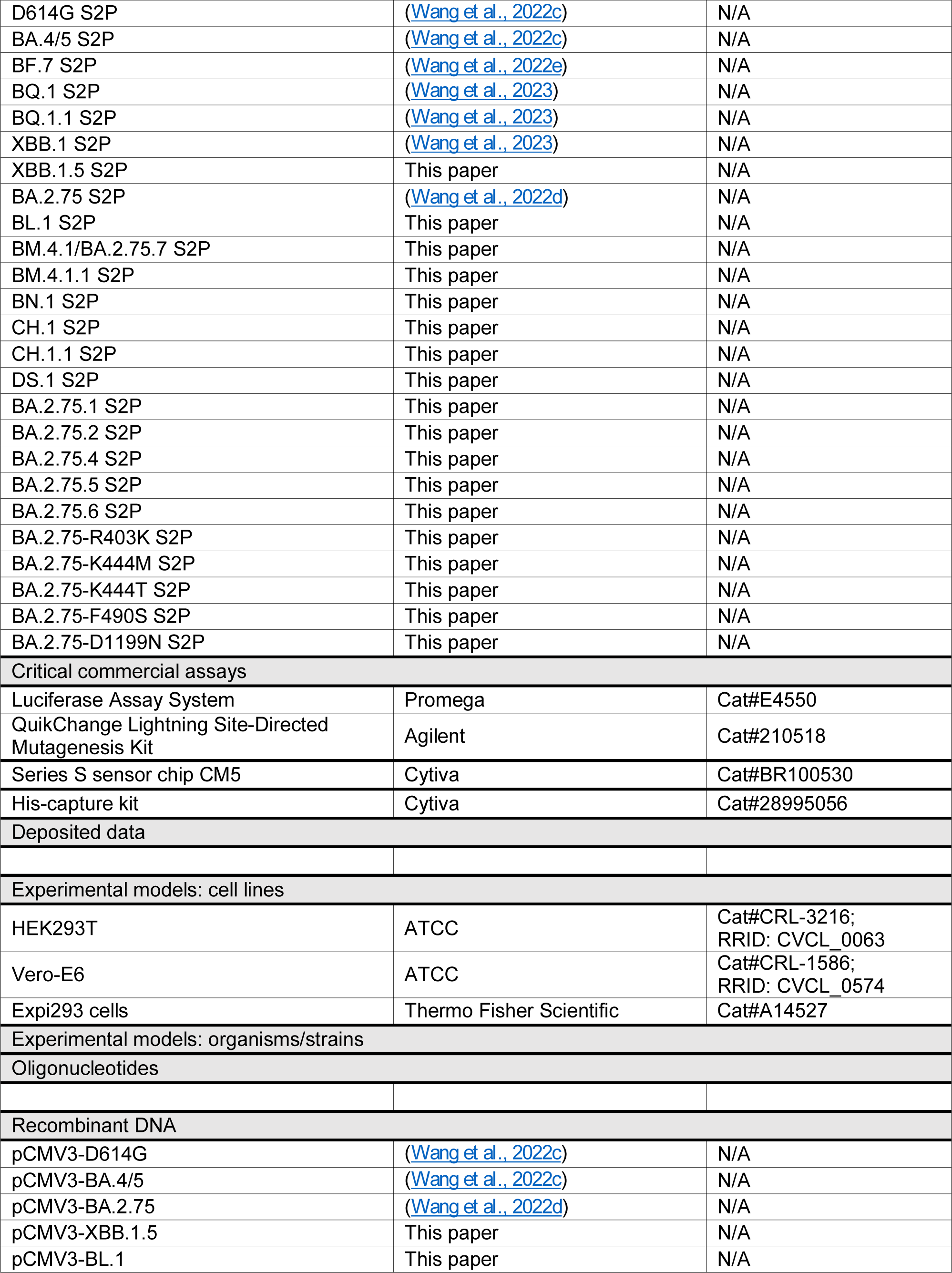
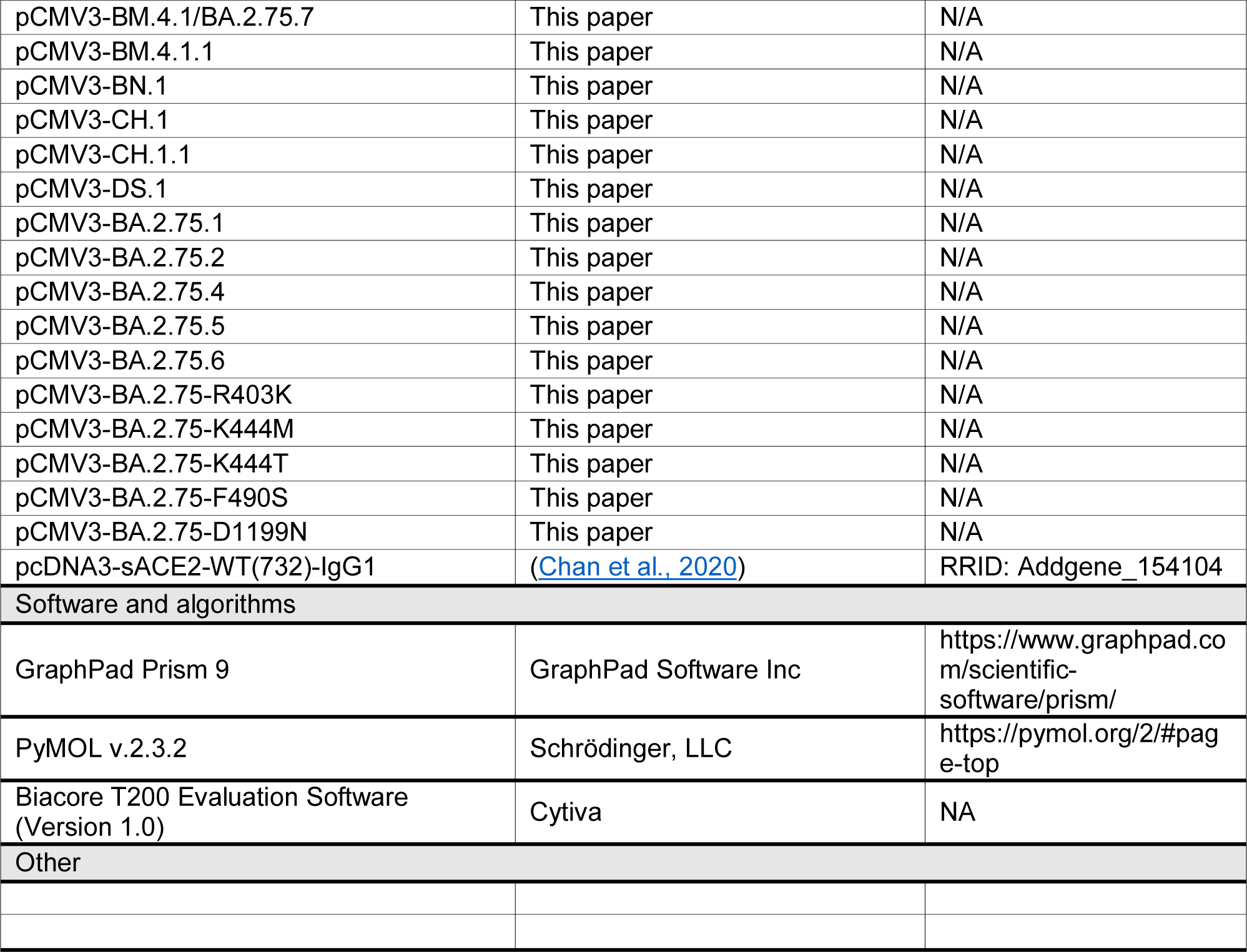

| REAGENT or RESOURCE | SOURCE | IDENTIFIER |
| --- | --- | --- |
| Antibodies |  |  |
| C1520 | (Wang et al., 2022h) | N/A |
| C1717 | (Wang et al., 2022h) | N/A |
| S3H3 | (Hong et al., 2022) | N/A |
| Brii-196 | (Ju et al., 2020) | N/A |
| Omi-3 | (Nutalai et al., 2022) | N/A |
| Omi-18 | (Nutalai et al., 2022) | N/A |
| BD-515 | (Cao et al., 2021) | N/A |
| COVOX-222 | (Dejnirattisai et al., 2021) | N/A |
| XGv051 | (Wang et al., 2022b) | N/A |
| XGv347 | (Wang et al., 2022a) | N/A |
| ZCB11 | (Zhou et al., 2022) | N/A |
| S2E12 | (Tortorici et al., 2020) | N/A |
| COV2-2196 | (Zost et al., 2020) | N/A |
| LY-CoV1404 | (Westendorf et al., 2022) | N/A |
| 2-7 | (Liu et al., 2020) | N/A |
| XGv289 | (Wang et al., 2022a) | N/A |
| XGv264 | (Wang et al., 2022b) | N/A |
| S309 | (Pinto et al., 2020) | N/A |
| P2G3 | (Fenwick et al., 2022) | N/A |
| SP1-77 | (Luo et al., 2022) | N/A |
| BD55-5840 | (Cao et al., 2022) | N/A |
| BD55-3152 | (Cao et al., 2022) | N/A |
| XGv282 | (Wang et al., 2022a) | N/A |
| BD-804 | (Du et al., 2021) | N/A |
| A19-46.1 | (Wang et al., 2021) | N/A |
| 35B5 | (Wang et al., 2022g) | N/A |
| JMB2002 | (Yin et al., 2022) | N/A |
| Brii-198 | (Ju et al., 2020) | N/A |
| COV2-2130 | (Zost et al., 2020) | N/A |
| 10-40 | (Liu et al., 2022) | N/A |
| Bacterial and virus strains |  |  |
| VSV-G pseudotyped ΔG-luciferase | Kerafast | Cat#EH1020-PM |
| Biological samples |  |  |
| 3 Shots WT sera | (Wang et al., 2023) | N/A |
| 3 Shots WT + Bivalent sera | (Wang et al., 2023) | N/A |
| BA.2 breakthrough sera | (Wang et al., 2023) | N/A |
| BA.4/5 breakthrough sera | (Wang et al., 2023) | N/A |
| Chemicals, peptides, and recombinant proteins |  |  |
| Polyethylenimine (PEI) | Polysciences Inc. | Cat#23966-100 |
| hACE2 | (Wang et al., 2022c) | N/A |
| D614G S2P | (Wang et al., 2022c) | N/A |
| BA.4/5 S2P | (Wang et al., 2022c) | N/A |
| BF.7 S2P | (Wang et al., 2022e) | N/A |
| BQ.1 S2P | (Wang et al., 2023) | N/A |
| BQ.1.1 S2P | (Wang et al., 2023) | N/A |
| XBB.1 S2P | (Wang et al., 2023) | N/A |
| XBB.1.5 S2P | This paper | N/A |
| BA.2.75 S2P | (Wang et al., 2022d) | N/A |
| BL.1 S2P | This paper | N/A |
| BM.4.1/BA.2.75.7 S2P | This paper | N/A |
| BM.4.1.1 S2P | This paper | N/A |
| BN.1 S2P | This paper | N/A |
| CH.1 S2P | This paper | N/A |
| CH.1.1 S2P | This paper | N/A |
| DS.1 S2P | This paper | N/A |
| BA.2.75.1 S2P | This paper | N/A |
| BA.2.75.2 S2P | This paper | N/A |
| BA.2.75.4 S2P | This paper | N/A |
| BA.2.75.5 S2P | This paper | N/A |
| BA.2.75.6 S2P | This paper | N/A |
| BA.2.75-R403K S2P | This paper | N/A |
| BA.2.75-K444M S2P | This paper | N/A |
| BA.2.75-K444T S2P | This paper | N/A |
| BA.2.75-F490S S2P | This paper | N/A |
| BA.2.75-D1199N S2P | This paper | N/A |
| Critical commercial assays |  |  |
| Luciferase Assay System | Promega | Cat#E4550 |
| QuikChange Lightning Site-Directed Mutagenesis Kit | Agilent | Cat#210518 |
| Series S sensor chip CM5 | Cytiva | Cat#BR100530 |
| His-capture kit | Cytiva | Cat#28995056 |
| Deposited data |  |  |
| Experimental models: cell lines |  |  |
| HEK293T | ATCC | Cat#CRL-3216;<br>RRID: CVCL_0063 |
| Vero-E6 | ATCC | Cat#CRL-1586;<br>RRID: CVCL_0574 |
| Expi293 cells | Thermo Fisher Scientific | Cat#A14527 |
| Experimental models: organisms/strains |  |  |
| Oligonucleotides |  |  |
| Recombinant DNA |  |  |
| pCMV3-D614G | (Wang et al., 2022c) | N/A |
| pCMV3-BA.4/5 | (Wang et al., 2022c) | N/A |
| pCMV3-BA.2.75 | (Wang et al., 2022d) | N/A |
| pCMV3-XBB.1.5 | This paper | N/A |
| pCMV3-BL.1 | This paper | N/A |
| pCMV3-BM.4.1/BA.2.75.7 | This paper | N/A |
| pCMV3-BM.4.1.1 | This paper | N/A |
| pCMV3-BN.1 | This paper | N/A |
| pCMV3-CH.1 | This paper | N/A |
| pCMV3-CH.1.1 | This paper | N/A |
| pCMV3-DS.1 | This paper | N/A |
| pCMV3-BA.2.75.1 | This paper | N/A |
| pCMV3-BA.2.75.2 | This paper | N/A |
| pCMV3-BA.2.75.4 | This paper | N/A |
| pCMV3-BA.2.75.5 | This paper | N/A |
| pCMV3-BA.2.75.6 | This paper | N/A |
| pCMV3-BA.2.75-R403K | This paper | N/A |
| pCMV3-BA.2.75-K444M | This paper | N/A |
| pCMV3-BA.2.75-K444T | This paper | N/A |
| pCMV3-BA.2.75-F490S | This paper | N/A |
| pCMV3-BA.2.75-D1199N | This paper | N/A |
| pcDNA3-sACE2-WT(732)-IgG1 | (Chan et al., 2020) | RRID: Addgene_154104 |
| Software and algorithms |  |  |
| GraphPad Prism 9 | GraphPad Software Inc | https://www.graphpad.com/scientific-software/prism/ |
| PyMOL v.2.3.2 | Schrödinger, LLC | https://pymol.org/2/#page-top |
| Biacore T200 Evaluation Software (Version 1.0) | Cytiva | NA |
| Other |  |  |

### Resource availability

#### Lead contact

Further information and requests for resources should be directed to and will be fulfilled by the lead contact, David D. Ho.

### Materials availability

All requests for resources and reagents should be directed to and will be fulfilled by the Lead Contact, David D. Ho. This includes selective cell lines, plasmids, antibodies, viruses, serum, and proteins. All reagents will be made available on request after completion of a Material Transfer Agreement.

### Data and code availability

**• Data**

Data reported in this paper will be shared by the lead contact upon request.

**•** Code

This paper does not report original code.

**•** All other items

Any additional information required to reanalyze the data reported in this paper is available from the lead contact upon request.

### Experimental model and subjects

#### Sample collection

The sera samples were all collected at Columbia University Irving Medical Center or at the University of Michigan through the Immunity-Associated with SARS-CoV-2 Study (IASO), and the collections were conducted under protocols reviewed and approved by the Institutional Review Board of Columbia University or the Institutional Review Board of the University of Michigan Medical School (Simon et al., 2022). All subjects provided written informed consent. Sera from individuals who received three doses of either the mRNA-1273 or BNT162b2 vaccines are described in the text as “3 shots WT”. Sera from individuals who received three doses of either the mRNA-1273 or BNT162b2 vaccines followed by a bivalent mRNA vaccine are described in the text as “3 shots WT + bivalent”. Sera from individuals who were infected by an Omicron subvariant (BA.2) following vaccinations were collected from December 2021 to May 2022 and are described in the text as “BA.2 breakthrough”. Sera from individuals who received vaccinations and were subsequently infected by an Omicron subvariant (BA.4/5) were collected from July 2022 to August 2022 and are described in the text as “BA.4/5 breakthrough”. To confirm prior SARS-CoV-2 infection status, anti-nucleoprotein (NP) ELISA tests were performed on all serum samples, as well as DNA sequencing to determine the variant involved in breakthrough cases. Clinical information on the different cohorts of study subjects is provided in **Table S2**.

### Cell lines

Vero-E6 (CRL-1586) cells and HEK293T (CRL-3216) cells were obtained from the ATCC and cultured at 37°C with 5% CO_2_ in Dulbecco modified Eagle medium (DMEM) with 10% fetal bovine serum (FBS) and 1% penicillin-streptomycin. Expi293 (A14527) cells were bought from Thermo Fisher Scientific and maintained in Expi293 Expression Medium following the manufacturer’s instructions. Vero-E6 cells are from African green monkey kidneys. HEK293T cells and Expi293 cells are of female origin.

### Method details

#### Plasmids

The antibody sequences for the heavy chain variable (VH) and the light chain variable (VL) domains were synthesized by GenScript, and then cloned into the gWiz vector to produce antibody expression plasmids. For the packaging plasmids for pseudoviruses, mutations were made by using the QuikChange II XL site-directed mutagenesis kit (Agilent) on the BA.2.75 construct that we previously generated (Wang et al., 2022d). For the soluble spike expression plasmids, the 2P substitutions (K986P, V987P) and a “GSAS” substitution in the furin cleavage site (682-685aa) were introduced in the ectodomain (1-1208aa in WA1) of each of the spikes and then fused with a 8x His-tag at the C-terminus as previously described (Wrapp et al., 2020). All constructs were verified using Sanger sequencing prior to use.

### Protein expression and purification

The gWiz-antibody, paH-spike, or pcDNA3-sACE2-WT(732)-IgG1 (Addgene plasmid #154104) (Chan et al., 2020) plasmid was transfected into Expi293 cells using PEI at a ratio of 1:3, and then the supernatants were collected after five days. The antibodies and human ACE2 fused to a Fc tag were purified with Protein A Sepharose (Cytiva) following the manufacturer’s instructions. For SPR analysis, the human ACE2 protein was further purified with Superdex 200 Increase 10/300 GL column. Spike proteins were purified using Ni-NTA resin (Invitrogen) following the manufacturer’s instructions. The molecular weight and purity were checked by running the proteins on SDS-PAGE prior to use.

### Surface plasmon resonance (SPR)

The CM5 chip was immobilized with anti-His antibodies using the His Capture Kit (Cytiva) to capture the spike protein through the C-terminal His-tag. Serially diluted human ACE2-Fc protein was then flowed over the chip in HBS-EP+ buffer (Cytiva). Binding affinities were measured with the Biacore T200 system at 25 in the single-cycle mode. Data was analyzed by the Evaluation Software using the 1:1 binding model.

### Pseudovirus production

Pseudotyped SARS-CoV-2 (pseudoviruses) were produced in the vesicular stomatitis virus (VSV) background, in which the native VSV glycoprotein was replaced by SARS-CoV-2 and its variants, as previously described (Wang et al., 2022c). Briefly, plasmids containing the appropriate spike were transfected into HEK293T cells with PEI. After 24 hours, VSV-G pseudotyped ΔG-luciferase (G*ΔG-luciferase, Kerafast) was added, and then washed with culture medium three times before being cultured in fresh medium for another 24 hours. Pseudoviruses were then harvested, centrifuged, and then aliquoted and stored at −80 °C.

### Pseudovirus neutralization assay

Each SARS-CoV-2 pseudovirus was titrated before use in the neutralization assay. Serially diluted heat-inactivated sera or antibodies were added in 96-well plates, starting at 1:100 dilution for sera and 10 µg/mL for antibodies. Then, pseudoviruses were added and incubated at 37 °C for 1 hour. In each plate, wells containing only pseudoviruses were included as controls. Vero-E6 cells were then added at a density of 3 × 10^4^ cells per well and incubate at 37 °C for an additional 10 hours. Cells were lysed and luminescence was determined by the Luciferase Assay System (Promega) and SoftMax Pro v.7.0.2 (Molecular Devices) according to the manufacturers’ instructions. Data were analyzed in GraphPad Prism v.9.3.

### Phylogenetic analysis

The genome sequences for each BA.2.75 subvariants were obtained from GISAID database (Accession: EPI_ISL_14217529, EPI_ISL_16926267, EPI_ISL_15050799, EPI_ISL_14908101, EPI_ISL_14536676, EPI_ISL_16581575, EPI_ISL_16939789, EPI_ISL_14492159, EPI_ISL_15611014, EPI_ISL_14434640, EPI_ISL_16040351, EPI_ISL_16434652, EPI_ISL_16954277, EPI_ISL_14536591 and EPI_ISL_13521515) to build the phylogenetic tree. The sequences were aligned by Muscle v3.8.31, and the low-quality sequencing sites with ‘N’ or ‘-’ were removed. The Maximum Likelihood tree was built in MEGA11 by Tamura-Nei model with 500 of bootstrap replication.

### Antibody footprint and mutagenesis analysis

All the structures were downloaded from Protein Data Bank (7XIX (BA.2 Spike), 8ASY (BA.2.75 RBD with ACE2), 7WK9 (S3H3), 7UAR (C1717), 7UAP (C1520), 7ZF3 (Omi-3), 7ZFB (Omi-18), 7CDI (Brii-196), 7OR9 (COVOX-222), 7WED (XGv347), 7K45 (S2E12), 7SD5(10-40), 7XCO (S309), 7WRV (JMB2002), 7WRZ (BD55-5840), 7WR8 (BD55-3152), 7WM0 (35B5), 7WLC (XGv282), 7WE9 (XGv289), 7UPY (SP1-77), 7U0D (A19-46.1), 7QTK (P2G3), 7MMO (LY-CoV1404), 7LSS (2-7), 7EYA (BD-804)) for analysis. The interface residues of each antibody were obtained by running the InterfaceResidues script from PyMOLwiki in PyMOL, and the edge of these residues was defined as the footprint after checking manually in the structure. Mutagenesis analyses were conducted in PyMOL. All the structure analysis figures were generated in PyMOL v.2.3.2 (Schrödinger, LLC).

### Antigenic cartography

The antigenic map was generated using the Racmacs package (https://acorg.github.io/Racmacs/, version 1.1.35) in R with 2000 optimization steps, a dilution step size of zero, and the minimum column basis parameter set to “none”. All distances between virus and serum positions on the map were optimized so that distances correspond to the fold decrease in neutralizing ID_50_ titer, relative to the maximum titer for each serum. Each unit of distance in any direction in the antigenic map corresponds to a two-fold change in the ID_50_ titer.

## QUANTIFICATION AND STATISTICAL ANALYSIS

Neutralization ID_50_ and IC_50_ values were determined by fitting a five-parameter dose-response curve using GraphPad Prism v.9.3. Statistical significance was evaluated using two-tailed Wilcoxon matched-pairs signed-rank tests in GraphPad Prism v.9.3.

---

**Figure S1.**
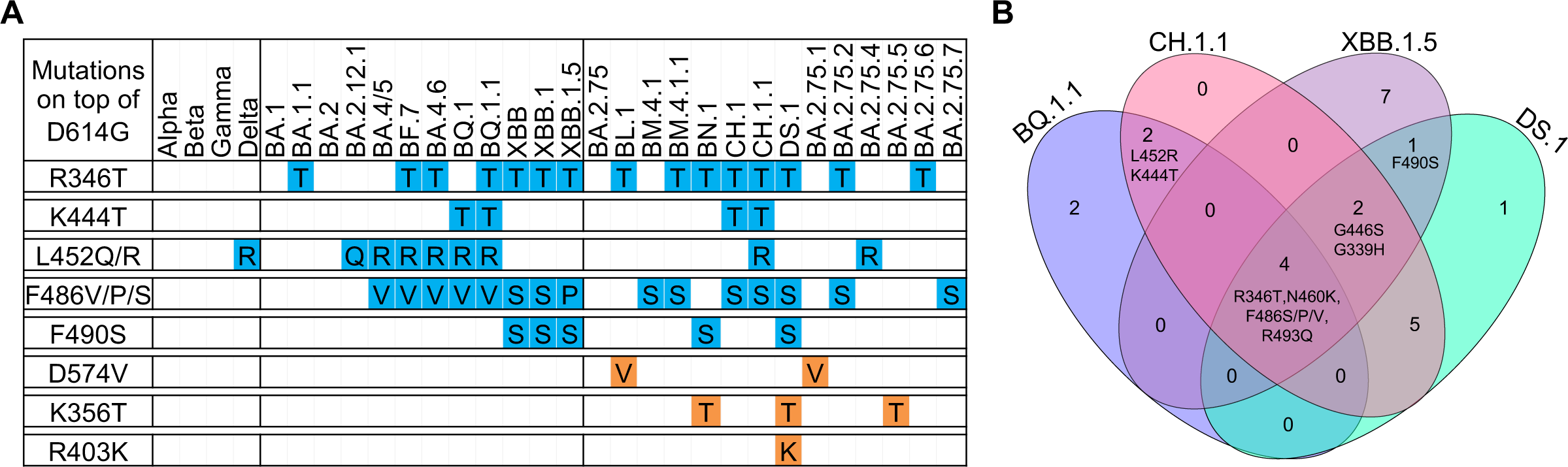
Key spike mutation in BA.2.75 subvariants. **(A)** Summary of spike mutations in SARS-CoV-2 VOCs. The convergent mutations are shown in blue and unique mutations among BA.2.75 subvariants are shown in orange. (B) The Venn diagram shows the number of shared spike mutations among BQ.1.1, XBB.1.5, CH.1.1, and DS.1 compared to BA.2. The RBD mutations shared by at least two subvariants are displayed. See also Figure 1.

**Figure S2.**
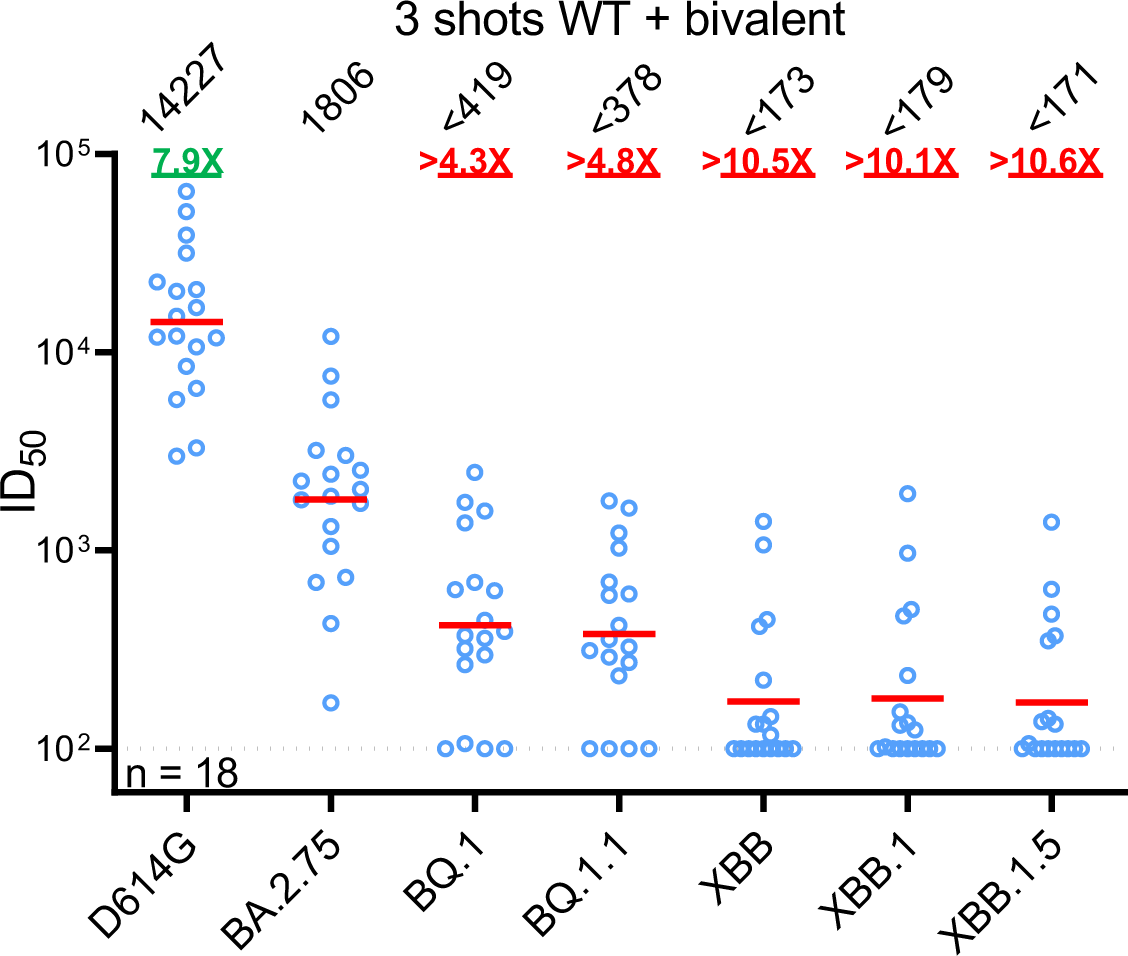
Neutralization of pseudotyped SARS-CoV-2 subvariants by polyclonal sera from “3 shots WT + bivalent” cohort. “3 shots WT + bivalent” refers to individuals vaccinated with three doses of the wildtype mRNA vaccine and subsequently one dose of a WA1/BA.5 bivalent mRNA vaccine. The results are shown as dots with geometric mean (red line). Values above the dots denote the raw geometric mean ID_50_ values and the sample size (n) is shown on the lower left. The limit of detection is 100 (dotted line). Comparisons were made against BA.2.75 and the fold changes in ID_50_ values are shown, with resistance to neutralization highlighted in red and sensitization in green. Statistically significant fold changes (*p* <0.05, determined by using two-tailed Wilcoxon matched-pairs signed-rank tests) are highlighted in bold. See also Figure 4.

**Table S1.**
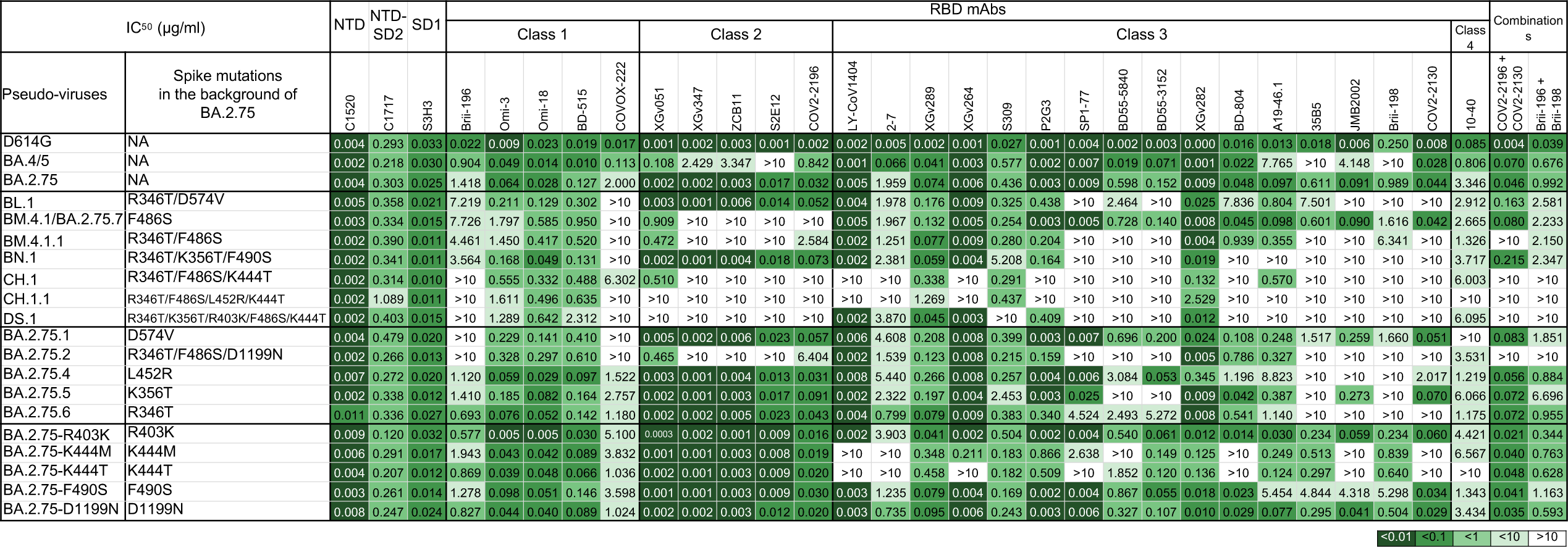
Neutralization IC_50_ values for indicated pseudotyped D614G, BA.4/5, BA.2.75, and BA.2.75 subvariants by mAbs. See also **Figure 3**.

**Table S2.** Demographics of clinical cohorts in this study. See also **Figure 4**.

| Sample ID | Vaccine type and infected strain | Days post-vaccination or<br>*infection (after last exposure) | Documented<br>COVID-19 | Age | Gender |
| --- | --- | --- | --- | --- | --- |
| 3 shots WT |  |  |  |  |  |
| 1 | mRNA-1273/mRNA-1273/mRNA-1273 | 29 | No | 66 | Female |
| 2 | BNT162b2/BNT162b2/BNT162b2 | 30 | No | 68 | Male |
| 3 | BNT162b2/BNT162b2/BNT162b2 | 14 | No | 64 | Female |
| 4 | BNT162b2/BNT162b2/BNT162b2 | 34 | No | 55 | Male |
| 5 | BNT162b2/BNT162b2/BNT162b2 | 34 | No | 45 | Male |
| 6 | BNT162b2/BNT162b2/BNT162b2 | 15 | No | 48 | Female |
| 7 | BNT162b2/BNT162b2/BNT162b2 | 29 | No | 71 | Male |
| 8 | BNT162b2/BNT162b2/BNT162b2 | 87 | No | 66 | Female |
| 9 | BNT162b2/BNT162b2/BNT162b2 | 84 | No | 26 | Male |
| 10 | mRNA-1273/mRNA-1273/mRNA-1273 | 23 | No | 28 | Female |
| 11 | BNT162b2/BNT162b2/mRNA-1273 | 32 | No | 39 | Male |
| 3 shots WT + bivalent |  |  |  |  |  |
| 12 | BNT162b2/BNT162b2/BNT162b2/Moderna Bivalent | 24 | No | 38 | Female |
| 13 | BNT162b2/BNT162b2/BNT162b2/Moderna Bivalent | 27 | No | 42 | Female |
| 14 | mRNA-1273//mRNA-1273/mRNA-1273/Moderna Bivalent | 24 | No | 36 | Male |
| 15 | BNT162b2/BNT162b2/BNT162b2/Pfizer Bivalent | 25 | No | 37 | Female |
| 16 | BNT162b2/BNT162b2/BNT162b2/Pfizer Bivalent | 24 | No | 36 | Male |
| 17 | BNT162b2/BNT162b2/BNT162b2/Pfizer Bivalent | 25 | No | 49 | Female |
| 18 | BNT162b2/BNT162b2/BNT162b2/Moderna Bivalent | 25 | No | 37 | Female |
| 19 | BNT162b2/BNT162b2/BNT162b2/Pfizer Bivalent | 26 | No | 45 | Male |
| 20 | BNT162b2/BNT162b2/mRNA-1273/Moderna Bivalent | 26 | No | 43 | Female |
| 21 | mRNA-1273/mRNA-1273/mRNA-1273/Moderna Bivalent | 29 | No | 32 | Female |
| 22 | BNT162b2/BNT162b2/BNT162b2/Pfizer Bivalent | 26 | No | 43 | Female |
| 23 | BNT162b2/BNT162b2/mRNA-1273/Moderna Bivalent | 29 | No | 38 | Female |
| 24 | BNT162b2/BNT162b2/BNT162b2/Moderna Bivalent | 28 | No | 38 | Female |
| 25 | BNT162b2/BNT162b2/mRNA-1273/Moderna Bivalent | 27 | No | 36 | Female |
| 26 | BNT162b2/BNT162b2/BNT162b2/Moderna Bivalent | 30 | No | 24 | Female |
| 27 | mRNA-1273/mRNA-1273/mRNA-1273/Moderna Bivalent | 30 | No | 32 | Female |
| 28 | BNT162b2/BNT162b2/mRNA-1273/Moderna Bivalent | 23 | No | 39 | Male |
| 29 | BNT162b2/BNT162b2/BNT162b2/Pfizer Bivalent | 30 | No | 26 | Female |
| BA.2 breakthrough |  |  |  |  |  |
| 30 | BNT162b2/BNT162b2/BA.2 | *14 | Yes | 50 | Female |
| 31 | BNT162b2/BNT162b2/BNT162b2/Ad26.COV2.S/BA.2 | *22 | Yes | 69 | Male |
| 32 | BNT162b2/BNT162b2/mRNA-1273/BA.2 | *16 | Yes | 32 | Male |
| 33 | mRNA-1273/mRNA-1273/mRNA-1273/BA.2 | *14 | Yes | 34 | Male |
| 34 | BNT162b2/BNT162b2/mRNA-1273/BA.2 | *19 | Yes | 33 | Female |
| 35 | BNT162b2/BNT162b2/mRNA-1273/BA.2 | *18 | Yes | 29 | Female |
| 36 | BNT162b2/BNT162b2/monovalent boost/BA.2 | *122 | Yes | 22 | Male |
| 37 | mRNA-1273/mRNA-1273/monovalent boost/BA.2 | *164 | Yes | 30 | Female |
| 38 | BNT162b2/BNT162b2/monovalent boost/BA.2 | *94 | Yes | 30 | Female |
| BA.4/5 breakthrough |  |  |  |  |  |
| 39 | BNT162b2/BNT162b2/BNT162b2/BA.5 | *29 | Yes | 44 | Female |
| 40 | BNT162b2/BNT162b2/mRNA-1273/BA.5 | *29 | Yes | 36 | Female |
| 41 | BNT162b2/BNT162b2/BNT162b2/BNT162b2/BA.5 | *31 | Yes | 54 | Female |
| 42 | BNT162b2/BNT162b2/BNT162b2/BNT162b2/BA.5 | *28 | Yes | 69 | Male |
| 43 | BNT162b2/BNT162b2/BNT162b2/BNT162b2/BA.5 | *42 | Yes | 44 | Male |
| 44 | BNT162b2/BNT162b2/BNT162b2/BNT162b2/BA.5 | *28 | Yes | 41 | Female |
| 45 | BNT162b2/BNT162b2/BNT162b2/BNT162b2/BA.5 | *31 | Yes | 29 | Female |
| 46 | BNT162b2/BNT162b2/BNT162b2/BNT162b2/BA.5 | *29 | Yes | 48 | Female |
| 47 | BNT162b2/BNT162b2/BNT162b2/BNT162b2/BA.5 | *29 | Yes | 49 | Female |
| 48 | BNT162b2/BNT162b2/mRNA-1273/BNT162b2/BA.5 | *28 | Yes | 37 | Female |
| 49 | mRNA-1273/mRNA-1273/BNT162b2/BA.5.2.1 | *29 | Yes | 29 | Female |
| 50 | BNT162b2/BNT162b2/BNT162b2/BA.5 | *22 | Yes | 61 | Female |
| 51 | mRNA-1273/mRNA-1273/mRNA-1273/BA.5 | *15 | Yes | 28 | Female |
| 52 | mRNA-1273/mRNA-1273/mRNA-1273/BA.5 | *21 | Yes | 24 | Female |
| 53 | BNT162b2/BNT162b2/BNT162b2/BA.5 | *75 | Yes | 35 | Female |
| 54 | BNT162b2/BNT162b2/mRNA-1273/BA.5 | *63 | Yes | 46 | Female |
| 55 | BNT162b2/BNT162b2/BNT162b2/BA.5 | *28 | Yes | 55 | Male |
| 56 | BNT162b2/BNT162b2/BNT162b2/BA.5 | *17 | Yes | 57 | Female |

